# The capacity to produce hydrogen sulfide (H_2_S) via cysteine degradation is ubiquitous in the human gut microbiome

**DOI:** 10.1101/2021.03.24.436781

**Authors:** Domenick J. Braccia, Xiaofang Jiang, Mihai Pop, A. Brantley Hall

## Abstract

As one of the three mammalian gasotransmitters, hydrogen sulfide (H_2_S) plays a major role in maintaining physiological homeostasis. Endogenously produced H_2_S plays numerous beneficial roles including mediating vasodilation and conferring neuroprotection. Due to its high membrane permeability, exogenously produced H_2_S originating from the gut microbiota can also influence human physiology and is implicated in reducing intestinal mucosal integrity and potentiating genotoxicity and is therefore a potential target for therapeutic interventions. Gut microbial H_2_S production is often attributed to dissimilatory sulfate reducers such as *Desulfovibrio* and *Bilophila* species. However, an alternative source for H_2_S production, cysteine degradation, is present in gut microbes, but the genes responsible for cysteine degradation have not been systematically annotated in gut microbes. To better understand the potential for H_2_S production via cysteine degradation by the human gut microbiome, we performed a comprehensive search for genes encoding cysteine-degrading genes in 4,644 bacterial genomes from the Unified Human Gastrointestinal Genome (UHGG) catalogue. We identified 407 gut bacterial species as putative cysteine degrading bacteria, 328 of which have not been previously implicated in H_2_S production. We identified the presence of at least one putative cysteine degrading bacteria in metagenomic data of 100% of 6,644 healthy subjects and the expression of cysteine-degrading genes in metatranscriptomics data of 100% of 59 samples. Additionally, putative cysteine-degrading bacteria are more abundant than sulfate reducing bacteria (p<2.2e-16). Overall, this study improves our understanding of the capacity for H_2_S production by the human gut microbiome and may help to inform interventions to therapeutically modulate gut microbial H_2_S production.

## Introduction

Hydrogen sulfide (H_2_S) is a consequential molecule produced by the gut microbiota with pleiotropic effects on human physiology. It is one of the three physiological gasotransmitters, along with carbon monoxide and nitric oxide, and is produced endogenously in many tissues including, but not limited to, the brain, heart and liver (1). Endogenous H_2_S production occurs via the enzymes cystathionine beta-synthase (CBS), cystathionine gamma-lyase (CSE) and 3-mercaptopyruvate sulfur transferase (MST) (2). CBS, CSE and MST are tightly regulated pyridoxal-5’-phosphate (PLP)-dependent enzymes and produce H_2_S primarily from the degradation of cysteine (3) (Figure 1B). H_2_S produced by these enzymes plays a litany of physiological roles including: suppression of oxidative stress in the brain, regulation of blood pressure through vasodilation and protection of hepatic stellate cells from cirrhosis in the liver (4). As a result, abnormally low endogenous levels of H_2_S are hypothesized to be an underlying cause of peripheral artery disease, and efforts have been made to measure serum levels of H_2_S quickly and non-invasively as a proxy for early detection of peripheral artery disease (5).

**Figure 1.**
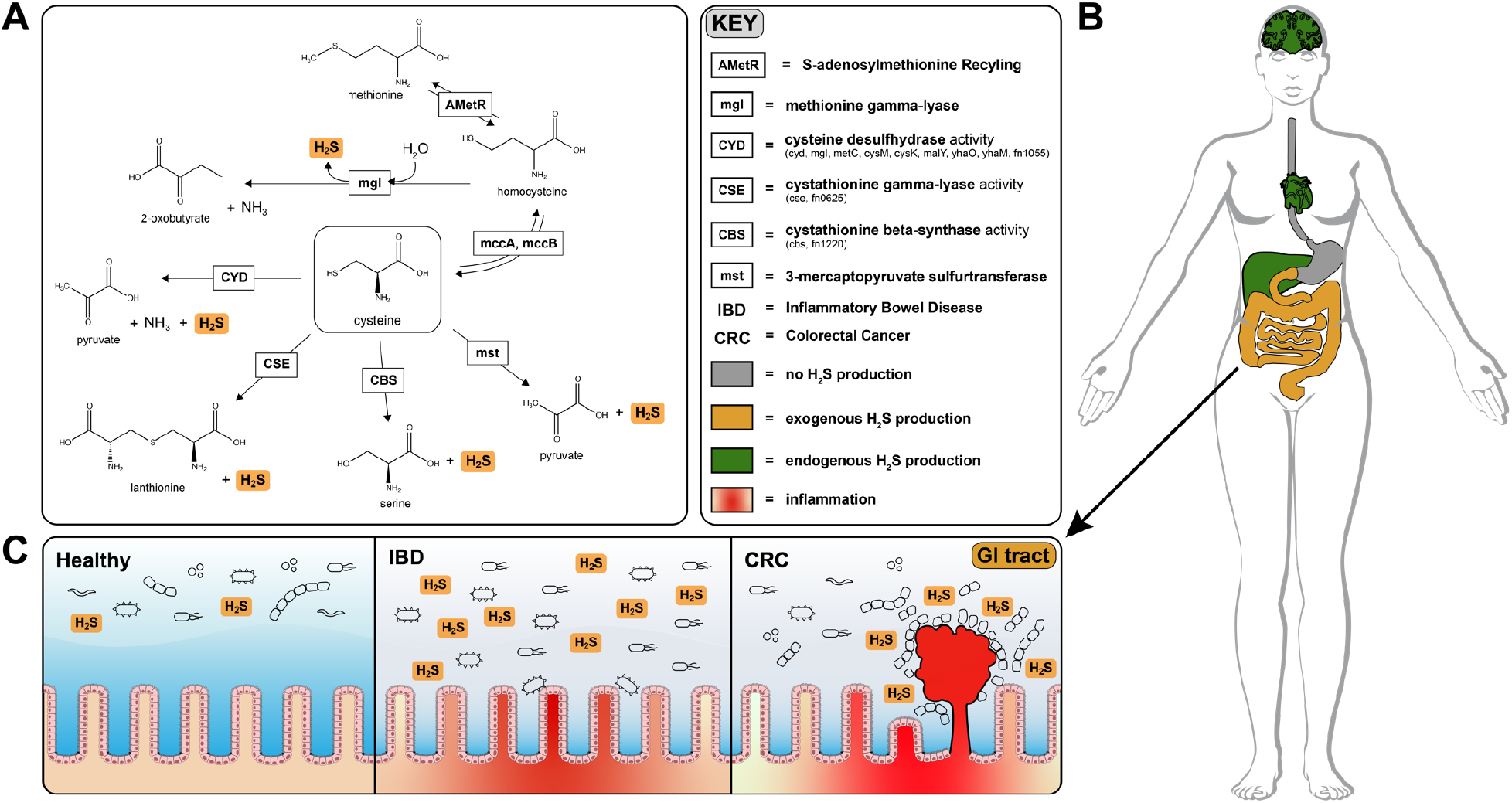
H_2_S production via cysteine degradation in the human gut microbiome. (A)Pathways of H_2_S production via cysteine degradation in the human gut microbiome. Pathways with labels ending in “activity” refer to a set of genes that convert cysteine to the same products. Genes involved in: CYD = (cyd, mgl, metC, cysM, cysK, malY, yhaO, yhaM, fn1055); CSE = (cse, fn0625); CBS = (CBS, fn1220). AMetR refers to the process of AdoMet recycling present in *Bacillus subtilis* involving genes mtnN, luxS and various methylases (45).(B) Visualization of H_2_S production across human tissues. H_2_S is produced endogenously in the brain, liver and heart via cysteine degradation and is tightly regulated to avoid toxic effects of H_2_S overproduction. Refer to the KEY for descriptions of organ color coding. (C) Physiological effects of H_2_S on the gut. Emphasis is placed on the difference between healthy versus IBD and CRC. In the IBD gut, H_2_S is thought to contribute to the degradation of the protective mucosal barrier which could cause or exacerbate inflammation. In CRC, various *Fusobacterium* species are closely associated with colonic tumors and are known H_2_S producers.

Microbes in the gastrointestinal tract also contribute to H_2_S production in humans. A majority of the microbially produced H_2_S originates in the colon, where estimates of luminal concentrations of hydrogen sulfide range from 0.3 mM to 3.4mM (6). The serum concentration of H_2_S in healthy individuals is difficult to measure but is estimated to range from 34.0 to 36.4 μM (7). H_2_S readily diffuses across the intestinal epithelium and can enter circulation influencing host physiology (8). Excessive production of H_2_S by gut microbes has been linked with decreased mucosal integrity through reduction of mucosal disulfide bonds (9), inhibition of colonocyte butyrate oxidation via cytochrome-c inhibition (10), and genotoxicity (8) (Figure 1C).

While the mammalian pathways of H_2_S production have been well studied, the contribution of gut-microbial H_2_S production to circulating H_2_S levels and the subsequent systemic effects on human physiology are largely unknown. The first step towards a better understanding of the effects of H_2_S on human physiology is to identify which microbial species are responsible for H_2_S production. There are two major sources for H_2_S production in the human gut microbiota, dissimilatory sulfate reduction and the degradation of the sulfur-containing amino acids cysteine and methionine (11). We must note that sulfate is first reduced to sulfite before H_2_S is produced, however, we refer to this process as sulfate reduction for the remainder of this work.

In the literature, H_2_S production is often attributed to the well-characterized dissimilatory sulfate reduction pathway (4). Common representatives of sulfate reducing bacteria (SRB) are found in the class *Deltaproteobacteria* with *Desulfovibrio* spp. and *Bilophila wadsworthia* being the most abundant representatives in the human gut (10). Sulfate and sulfite are used by SRB as terminal electron acceptors for anaerobic respiration (12). While SRB are prevalent in human populations, their relative abundances are generally very low and are dependent on ecological interactions with other hydrogenotrophs, such as methanogens and acetogens (10,13,14).

Unlike the comprehensively-characterized pathways for dissimilatory sulfate reduction, the species of the gut microbiome responsible for H_2_S production via degradation of sulfur-containing amino acids have not been comprehensively characterized. Gut microbial involvement in amino acid fermentation has garnered recent attention, as many physiologically relevant downstream metabolites are produced by gut microbial degradation of amino acids (15) (Figure 1A). Depending on dietary intake, a pool of sulfur-containing amino acids is available for fermentation by gut microbiota (16). Recent studies have demonstrated that cysteine supplementation leads to far more H_2_S production than inorganic sulfate supplementation underscoring the comparative importance of the cysteine-degradation pathway in total H_2_S production (12–14).

Methane (CH_4_) is primarily produced by the methanogen *Methanobrevibacter smithii* (17) and is one of the primary gases present in mammalian flatus. Sulfate-reducing bacteria and methanogens have been historically considered mutually exclusive in microbial communities due to the competition for hydrogen (10). However, experiments carried out on human flatus have shown that both H_2_S and CH_4_ production occurs simultaneously in some individuals, seemingly contradicting the notion that methanogens and sulfate-reducing bacteria cannot co-exist (6).

It is important to delineate between H_2_S produced via dissimilatory sulfate reduction and H_2_S produced via cysteine degradation because different approaches are necessary to modulate these two sources of H_2_S production. Because of the poor annotation of the genes which produce H_2_S via cysteine degradation across species of the gut microbiome, the relative contributions of H_2_S production are unclear. To address this gap, we designed a bioinformatic approach to first identify putative cysteine-degrading bacteria in the human gut microbiome and then compare the relative abundances of putative cysteine-degrading bacteria and sulfate-reducing bacteria across metagenomic data from inflammatory bowel disease, colorectal cancer and healthy cohorts.

## Results

### Cysteine-degrading genes are widely distributed in and expressed by the human gut microbiome

To identify species capable of H_2_S production via cysteine degradation in the human gut microbiome, we curated a list of enzymes experimentally proven to produce H_2_S, and searched for these enzymes across 4,644 species in the Unified Human Gastrointestinal Genome (UHGG) collection (18) (Figure 2, Supplementary Table 1). This collection comprises 4,644 non-redundant genome sequences from species representatives generated by clustering 204,938 genome sequences from bacteria known to inhabit the human gut.

**Figure 2.**
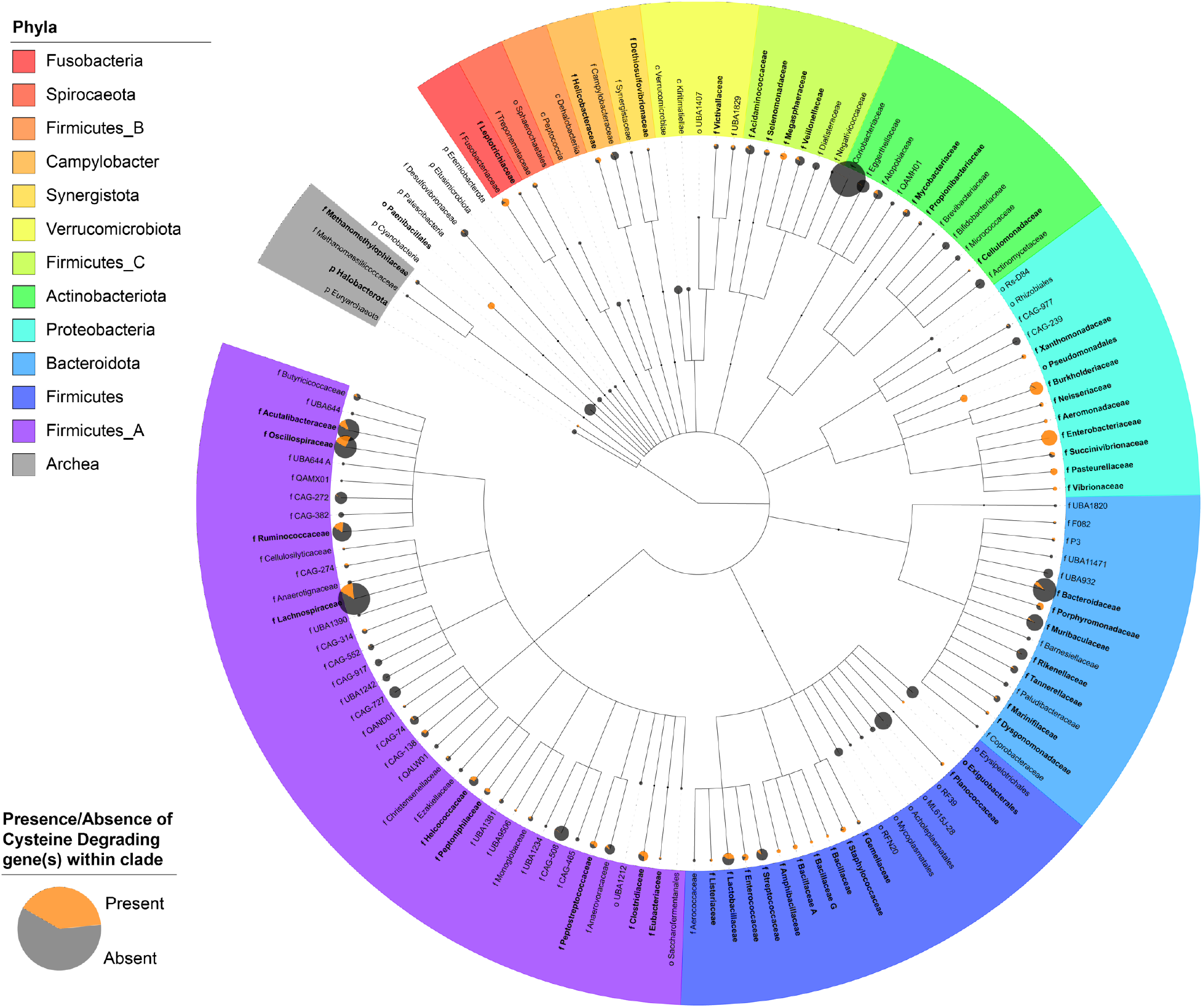
Distribution of putative cysteine-degrading bacteria across the United Human Gastrointestinal Genome (UHGG) collection. A taxonomic tree showing the distribution of putative cysteine-degrading bacteria across the 4,644 genomes of the representative UHGG collection. Leaves of the tree are shown to the family level and only genera with >= 3 subspecies are included. Phyla are labeled by color and pie charts at the leaf nodes correspond to presence or absence of cysteine-degrading genes whose expression results in H_2_S production. The relative size of the pie chart represents the number of species in the family shown.

Of the representative UHGG species, 18.4% (855/4,644) contain one or more cysteine-degrading genes (Figure 3A) whereas just 0.6% (27/4,644) contain the sulfate-reducing genes *dsrAB*. Aside from known cysteine-degrading bacterial species compiled in the manual curation step, an additional 406 previously cultured species were found to contain one or more cysteine-degrading genes (Figure 2, names in bold). Furthermore, 10.8% (44/406) of these species have evidence of H_2_S production, 8.6% (35/406) showed no signs of H_2_S production, and 80.8% (328/406) have had no prior test for H_2_S production (Supplementary Table 1). Additionally, 550 metagenome-assembled genomes (MAGs) were found to contain one or more cysteine-degrading genes. No UHGG genomes contain both *dsrAB* and a cysteine-degrading gene, while many genomes contain multiple cysteine-degrading genes (Figure 3A).

**Figure 3.**
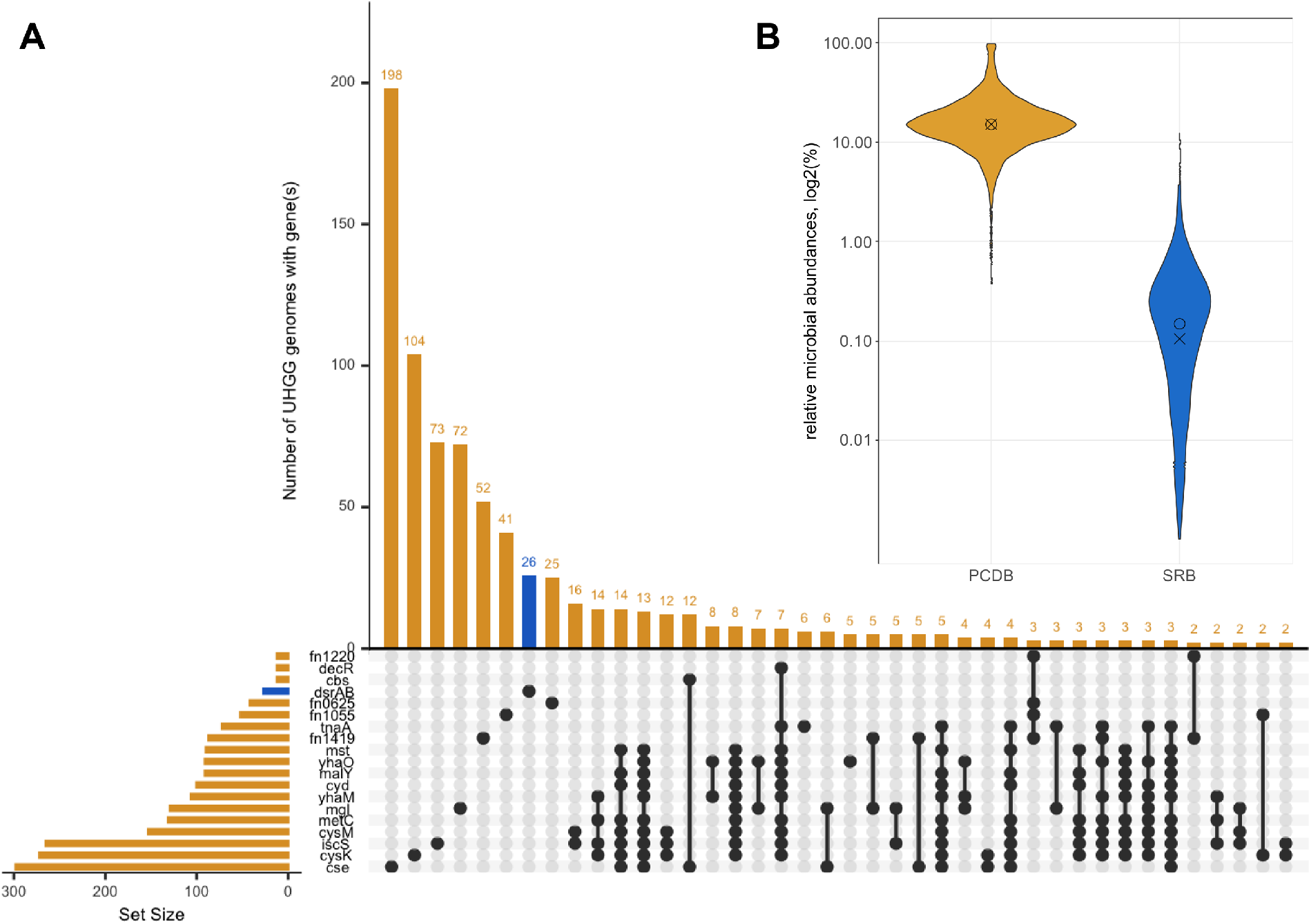
Comparing putative cysteine-degrading bacteria (PCDB) to sulfate reducing bacteria (SRB). Color key: orange = cysteine-degrading genes and bacteria containing such genes; blue = sulfate reducing genes and bacteria containing such genes. (A) Overlap of gene hits in the UHGG collection. The y-axis shows the number of genomes receiving one or more hits to H_2_S producing genes CYD activity genes = (cyd, metC, cysM, cysK, malY, yhaO, yhaM, fn1055); CSE activity genes = (cse, fn0625); CBS activity genes = (cbs, fn1220); sulfate reducing genes = (dsrAB). The x-axis shows genes which co-occur in UHGG genomes. For example, the gene cse appeared in 198 genomes individually and appeared alongside the gene cbs in 12 genomes. The dissimilatory sulfate reducing gene dsrAB occurred in only 27 genomes, and did not co-occur with any cysteine-degrading genes searched. All species receiving at least one hit to a cysteine-degrading gene are considered putative cysteine-degrading bacteria. (B) Relative abundance of putative cysteine-degrading bacteria and sulfate reducing bacteria among 6,644 healthy controls provided by curatedMetagenomicData (19) (p<2.2×10^−16^, two-sided Wilcoxon rank sum test).

### Widespread potential for H_2_S production via cysteine degradation in the human gut microbiome

The prevalence and relative abundance of putative cysteine-degrading bacteria and sulfate-reducing bacteria was calculated for 10,700 metagenomic samples from healthy, inflammatory bowel disease, colorectal cancer and adenoma cohorts (19–22). Among the 6,644 healthy subjects, there is a markedly higher (W = 44,133,484; p < 2.2e-16, two-sided Wilcoxon rank sum test) relative abundance of putative cysteine-degrading bacteria compared to sulfate-reducing bacteria suggesting that cysteine degradation contributes considerably to H_2_S production for the average healthy person (Figure 3B). Cysteine degrading genes are also widespread in healthy populations with 100% of the 6,644 healthy subjects containing at least one putative cysteine-degrading bacteria in their gut microbiome.

To confirm the transcription of cysteine-degrading genes and sulfate-reducing genes, we analyzed metatranscriptomic data from a dietary intervention study by Lawrence et al. (23). The study relied on stool sample collections before and after plant-based and animal-based diet interventions to evaluate the effects of diet on microbial gene regulation and community composition. Our analysis revealed that 100% of samples from this study (59/59) showed non-trivial expression (RPKM >= 1) of one or more cysteine-degrading genes. While 61% of samples (36/59) contained expression of both dissimilatory sulfate reduction and cysteine degradation, 36% of samples (21/59) had cysteine degradation genes as the sole source of H_2_S production (Figure S4). Methionine gamma-lyase is the most actively transcribed H_2_S producing gene, and 3-mercaptopyruvate sulfurtransferase is generally the least transcribed (Figure S2). The primary dissimilatory sulfate reductase genes, *dsrAB*, appear constitutively expressed across all three diets. Comparatively, expression of cysteine-degrading genes appears to be more sporadic across diet conditions and slightly lower in both plant and animal-based diets compared to baseline. These results suggest that for more than one third of individuals cysteine degradation may be the dominant pathway for H_2_S production.

### Core genes from dissimilatory sulfate reduction and methanogenesis are co-expressed

Though *in vitro* assays have indicated that methanogens and sulfate-reducing bacteria compete for hydrogen and may thus mutually exclude one another (10), core genes involved in dissimilatory sulfate reduction (Figure S2) and methanogenesis (Figure S3) are simultaneously expressed in 8% (5/59) of samples obtained from healthy individuals (Figure S4). Additionally, one or more cysteine-degrading genes were expressed simultaneously with methanogenesis genes in 10% (6/59) of samples.

### Increased relative abundance of H_2_S producing bacteria in the colorectal cancer gut microbiome

Next, we assessed the relative abundance of putative cysteine-degrading bacteria and sulfate-reducing bacteria in individuals with the two most common clinical manifestations of inflammatory bowel disease (IBD), Crohn’s disease and ulcerative colitis, colorectal cancer (CRC) and healthy controls. Putative cysteine-degrading bacteria are significantly more abundant than sulfate-reducing bacteria across IBD and CRC populations from metagenomic samples derived from curatedMetagenomicData (19), the Human Microbiome Project 2 (HMP2) (20), PRISM (21) and Lewis *et al*. (22) (all p < 2.2×10^−16^) (Figure 4A-D).

**Figure 4.**
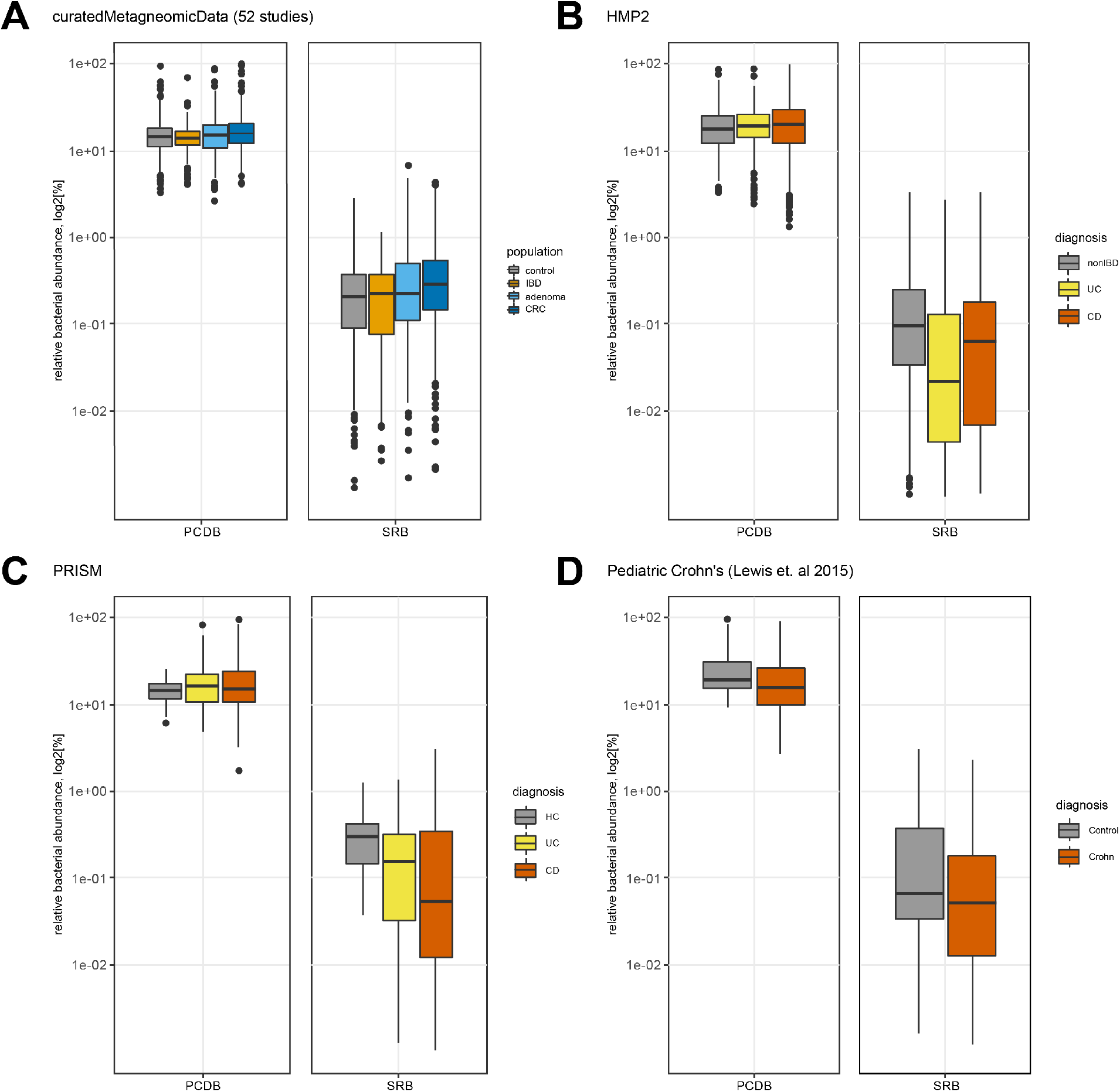
Putative cysteine-degrading bacteria (PCDB) are more prevalent than sulfate-reducing bacteria (SRB) among individuals with IBD, CRC, adenoma and healthy controls. Log2-transformed relative abundances of putative cysteine-degrading bacteria and sulfate-reducing bacteria across healthy and diseased populations. Relative abundances were calculated using Kraken2 (40) (see methods section). (A) Data obtained from curatedMetagenomicData (19). Number of samples per disease category: control = 560, CRC = 352, adenoma = 143, IBD = 148. (B) Data obtained from HMP2 (20). Number of samples per disease category: nonIBD = 359, ulcerative colitis (UC) = 367, Crohn’s disease (CD) = 591. (C) Data obtained from PRISM (21). Number of samples per disease category: control = 56, UC = 76, CD = 88. (D) Data obtained from Lewis et al. 2015 (22). Number of samples per disease category: control = 26, CD = 86.

Both putative cysteine-degrading bacteria and sulfate-reducing bacteria are significantly more abundant in CRC than in the respective control groups (Figure 4A). The strength of the association is similar for putative cysteine-degrading bacteria (W = 114,615; p < 3.4×10^−5^) and sulfate-reducing bacteria (W = 118,888, p < 1.5×10^−7^). In adenomas sulfate-reducing bacteria were found to be moderately differentially abundant (W = 44,330; p = 0.048) while putative cysteine-degrading bacteria were not found to be significantly differentially abundant (W = 38,494; p = 0.48).

The relative abundance of putative cysteine-degrading bacteria is moderately higher than controls for adults with Crohn’s disease in the HMP2 cohort, but not in the PRISM cohort. (HMP2: W = 114,116, p = 0.05; PRISM: W = 2,736, p = 0.27) and relatively similar for infants with Crohn’s disease (W = 862, p = 0.08). Likewise, putative cysteine-degrading bacteria are seen at similar relative abundance in adults with ulcerative colitis compared to healthy controls (HMP2: W = 70,404, p = 0.11; PRISM: W = 2,407, p = 0.20).

The relative abundance of sulfate-reducing bacteria tends to be lower in individuals with IBD compared to healthy controls. In adults with Crohn’s disease, there is a significantly lower relative abundance of sulfate-reducing bacteria compared to healthy controls (HMP2: W = 87,939, p = 1.0×10^−5^; PRISM: W = 1,435, p = 2.5×10^−5^). Infants with Crohn’s disease do not have a significant difference in relative abundance of sulfate-reducing bacteria (W = 964, p = 0.29) compared to healthy controls. Adults with ulcerative colitis have markedly lower relative abundance of sulfate-reducing bacteria compared to healthy controls (HMP2: W = 45,423, p = 4.5×10^−13^; PRISM: W = 1,382, p = 6.0×10^−4^).

## Discussion

Due to its role as a mammalian gasotransmitter, H_2_S plays important roles in maintaining physiological homeostasis. However, H_2_S may also cause deleterious effects in a concentration-dependent manner. Therefore, it is of great importance to understand the sources of exogenous H_2_S production in the gut in order to tease out the links between H_2_S and human physiology.

The source of gut microbial H_2_S production is often attributed to dissimilatory sulfate reduction, with far less attention given to H_2_S production via the degradation of the sulfur-containing amino acid cysteine. In fact, there has not been a microbiome-wide annotation of the potential for H_2_S production via cysteine degradation. The systematic annotation we performed in this study expands our understanding of which species can produce H_2_S in the gut, many of which have not been previously reported to have the capability for H_2_S production (Supplementary Table 1). Our analysis of shotgun sequenced metagenomic data from 6,644 metagenomic samples revealed that putative cysteine-degrading bacteria are ubiquitous inhabitants of the human gut microbiome and have significant higher relative abundance than sulfate-reducing bacteria. Furthermore, analysis of metatranscriptomic data demonstrates that cysteine-degrading genes are in fact expressed in the gut. These results suggest that cysteine degradation is likely a major source of microbial H_2_S production and may be the sole source of microbially produced H_2_S in some individuals. Therefore, cysteine degradation is an important aspect to consider when designing studies to assess the effects of H_2_S on human health or modulate gut microbial H_2_S production.

We also explored the relative abundance of putative cysteine-degrading bacteria in IBD and CRC to understand whether these bacteria could contribute to, or promote disease progression. We found that putative cysteine-degrading bacteria are significantly more abundant in CRC samples than in healthy controls. While sulfate-reducing bacteria are also increased in CRC compared to healthy controls, putative cysteine-degrading bacteria are far more abundant. This finding corroborates previous studies linking H_2_S and the progression of CRC (24) and highlights the need to identify the dominant source of H_2_S in the CRC gut. Putative cysteine-degrading bacteria were not differentially abundant between samples from IBD patients and healthy individuals but are more abundant than sulfate-reducing bacteria. Importantly, it still remains to be elucidated whether or not this difference in relative abundance directly translates to higher production of H_2_S via cysteine degradation in comparison with sulfate reduction.

Prior studies suggested that methanogens and sulfate-reducing bacteria are mutually exclusive, potentially due to their competition for hydrogen. These experiments did not consider cysteine degradation as a potential source of H_2_S. However, subsequent studies have reported the presence of both CH_4_ and H_2_S in the human flatus (6), seemingly contradicting this notion of mutual exclusivity of CH_4_ and H_2_S producing bacteria. To resolve this discrepancy, we examined the transcriptional co-occurrence of methanogens, sulfate-reducing bacteria, and

cysteine-degrading bacteria in the human gut and found the co-occurrence of all three pathways. This discrepancy between *in vitro* experiments and *in vivo* observations could be explained by the complex biogeography of the gut in which methanogens and sulfate-reducing bacteria occupy distinct niches or from H_2_S production via cysteine degradation.

There are many reactions in which H_2_S is formed as an intermediate, such as assimilatory sulfate reduction, however, these reactions do not result in significant production of H_2_S and are thus not relevant to total H_2_S production by the gut microbiome. Therefore, we limited our search for H_2_S producing bacteria to pathways in which H_2_S was the endpoint, or byproduct, and not just an intermediate of the pathway. Our search identified the genes for dissimilatory sulfate reduction in *Eggerthella* and *Gordinobacter* species. We have included these species as sulfate-reducing bacteria though there is little evidence to suggest that these species are true sulfate reducers (25,26). Further wet-lab validation of these claims is necessary to confirm *Eggerethella* spp. and *Gordinobacter* spp. as non-sulfate-reducing bacteria. We also note that our search for H_2_S producing genes included only the 4,644 representative genomes in UHGG. The full UHGG collection contains 204,938 non-redundant genomes with core and accessory gene information that may contain other putative H_2_S-producing sub-species that we did not analyze. Another potential shortcoming of this analysis is the overrepresentation of western countries in the data pool used. An expanded set of samples would be required to claim that putative-cysteine degrading bacteria are globally prevalent in the human gut microbiome.

Finally, we note that sulfate-reducing bacteria may be mucosally associated and present at low relative abundances which could mean that stool metagenomics may underestimate the true abundance of sulfate-reducing bacteria in the human gut.

In conclusion, we show that the relative abundance of putative cysteine-degrading bacteria is significantly higher than sulfate-reducing bacteria across healthy individuals as well as individuals with colorectal cancer and inflammatory bowel disease. These results bolster previous studies suggesting the importance of dietary cysteine in gut microbial H_2_S production. We also provide a comprehensive overview of putative cysteine-degrading bacteria complete with experimental evidence, or lack thereof, for H_2_S production in numerous experimental contexts. The systematic annotation of putative H_2_S-producing species performed in this study can serve as a resource for future studies examining the links between H_2_S and disease and could help these studies to tease out the concentration-dependent effects of H_2_S on human health. Overall, this work informs future approaches to modulate gut microbial H_2_S production via dietary interventions and may lead to an improved understanding of the complex interplay between H_2_S and human health and disease.

## Methods

### Curation of cysteine-degrading and sulfate-reducing genes

A search for potential hydrogen sulfide producing bacteria was conducted using *a priori* knowledge of dissimilatory sulfate reduction and sulfur-containing amino acid degradation by gut microbes. First, amino acid sequences of enzymes responsible for H_2_S production (11,27–31) were downloaded using the UniProtKB (32) web browser (www.uniprot.org). Links to UniProt entries used in the search space are listed in Supplementary Table 1 under the “search_space” sheet.

### Search for putative H2S producing bacteria in the human gut

Note that all BLAST tools used in this work are from the `blast+` command line package, version 2.8.1 (33). A protein BLAST database was created using the protein sequences as input to `makeblastdb` with option `-dbtype prot`. These protein sequences were then searched against 4,644 genome sequences from UHGG (18) using `blastp` (34). Hits were filtered based on two fairly conservative thresholds: E-value < 1×10^−110^ and amino acid identity > 50%. Then, depending on the nature of the sequence matches, the bacterial genomes receiving hits to H_2_S producing genes were labeled as either putative cysteine-degrading bacteria or sulfate-reducing bacteria. For instance, any hit to one or more of the cysteine-degrading genes was enough to consider the species to be a putative cysteine-degrading bacterium. However, in order to be considered capable of dissimilatory sulfate reduction, the genome must have received a hit for both *dsrA* and *dsrB* as they are subunits of the final functional protein. Putative cysteine-degrading bacteria across UHGG were then visualized by uploading a taxonomic tree in newick tree format to the iTOL (35) web interface (Figure 2). Pie charts in Figure 2 were generated by parsing the `blastp` output GFF files and uploading to the iTOL web interface. Figure 3A showing the overlap of gene hits to the UHGG collection was generated using UpSetR (36).

### Calculating relative abundances with Kraken2

The raw sequencing reads for the metagenomic samples used in this study were downloaded and extracted with NCBI’s SRA toolkit v2.10.9 (37). Quality control and adapter trimming of the fastq sequence files were done with the Trim Galore wrapper v0.6.6 (38). To remove potential human contaminants, quality-trimmed reads were screened against the human genome (hg19) with Bowtie2 v2.4.2 (39). Taxonomy profiling of the metagenomic cleaned reads were generated using Kraken2(2.0.8-beta) (40) to map against the pre-built database of the Unified Human Gastrointestinal Genome (UHGG) catalog (18).

### Analysis of putative sulfate-reducing bacteria

Amino acid sequences encoding dissimilatory sulfate reductase genes dsrA and dsrB were downloaded from UniProt (accession links in Supplementary Table 1). `blastp` was used to query 4,644 genomes from UHGG for additional species potentially performing dissimilatory sulfite reduction.

The search returned 27 valid hits (≥ 50% amino acid identity and E-value ≤ 1e-110 for both dsrA and dsrB) to bacteria under the phyla *Proteobacteria, Firmicutes*, and *Actinobacteria* (Supplementary Table 1, sheet labeled “sulfate_reduction_hits”). Hits to bacteria within the *Proteobacteria* phylum were expected, as the subphylum *Deltaproteobacteria* contains well-known sulfate-reducing bacteria. Hits to the *Firmicutes* species *Desulfitobacterium hafniense* were also expected since this taxon has been shown to reduce sulfite compounds to sulfide (41,42). Per Muller et al. (25), the presence of dsrAB in *Gordonibacter pamelaeae* (phylum *Actionobacteria*) is likely due to a lateral gene transfer event from the genus *Desulfitobacterium* as evidenced by the incongruence between phylogenies built using 16S rRNA gene sequencing and dsrAB gene sequencing. We replicated this phenomenon by constructing a phylogenetic tree from 27 sequences that match dsrAB within UHGG genomes (Figure S2). To construct the tree, a multiple sequence alignment of the 27 sequences using `mafft` version 7.307 with options `--maxiterate 1000` and `--localpair` was fed to `FastTree` version 2.1.9 with options `-nt`. The tree was then vizualized using the iTOL web interface.

Aside from the expected cases, we decided to include hits to *Firmicutes* and *Actinobacteria* species without experimental evidence of H_2_S production via dissimilatory sulfate reduction. Our rationale for including these species is to stay consistent with our inclusion of cysteine-degrading bacteria lacking experimental evidence of H_2_S production from cysteine degradation. We did not add the “putative” descriptor to this group of bacteria because, unlike the putative cysteine-degrading bacteria we identified, the vast majority of the species that turned up in our results are experimentally validated sulfate-reducing bacteria.

### Transcriptomic analysis of H_2_S producing genes and CH_4_ producing genes

We sought to confirm the active expression of H_2_S producing genes and CH_4_ producing genes alongside the existing genomic evidence presented using data from David et al. 2014 (23). In this study, 10 participants had RNA sequencing performed on their stool samples before and after a plant-based and animal-based diet intervention. Subjects waited 6 days before switching to the next diet and getting a baseline reading. Confirming the expression of H_2_S producing genes involved the following steps: 1. Metadata for samples was downloaded from the SRA run selector https://trace-ncbi-nlm-nih-gov. 2. Raw sequencing data was downloaded using `fasterq-dump` from the SRA toolkit version 2.10.9 (37). 3. Manually curated H_2_S producing genes were given as input to `diamond makedb` (43). 4. Raw RNA-seq data were then aligned against the manually curated protein database using the `diamond blastx` command with options `-b42.0 -c1` for better performance. The raw counts of reads mapped per gene were normalized to RPKM values for downstream analysis. The threshold for considering an H_2_S gene “expressed” was RPKM >=1. A sample was said to be “methane producing” if >= 90% of the genes involved in the methanogenesis pathway recruited one or more read mapping. The results were then parsed with a custom shell script and visualized in Figure S3 using the R package ggplot (44).

## Acknowledgements

D.J.B. is supported by National Science Foundation-funded Research Traineeship (COMBINE), X.J. issupported by the Intramural Research Program of the NIH, National Library of Medicine. M.P. is supported by NIH grant R01-AI-100947. B.H. is supported by startup funding from the University of Maryland. The authors declare no conflict of interests.

## Supplementary Material

**Supplementary Figure 1.**
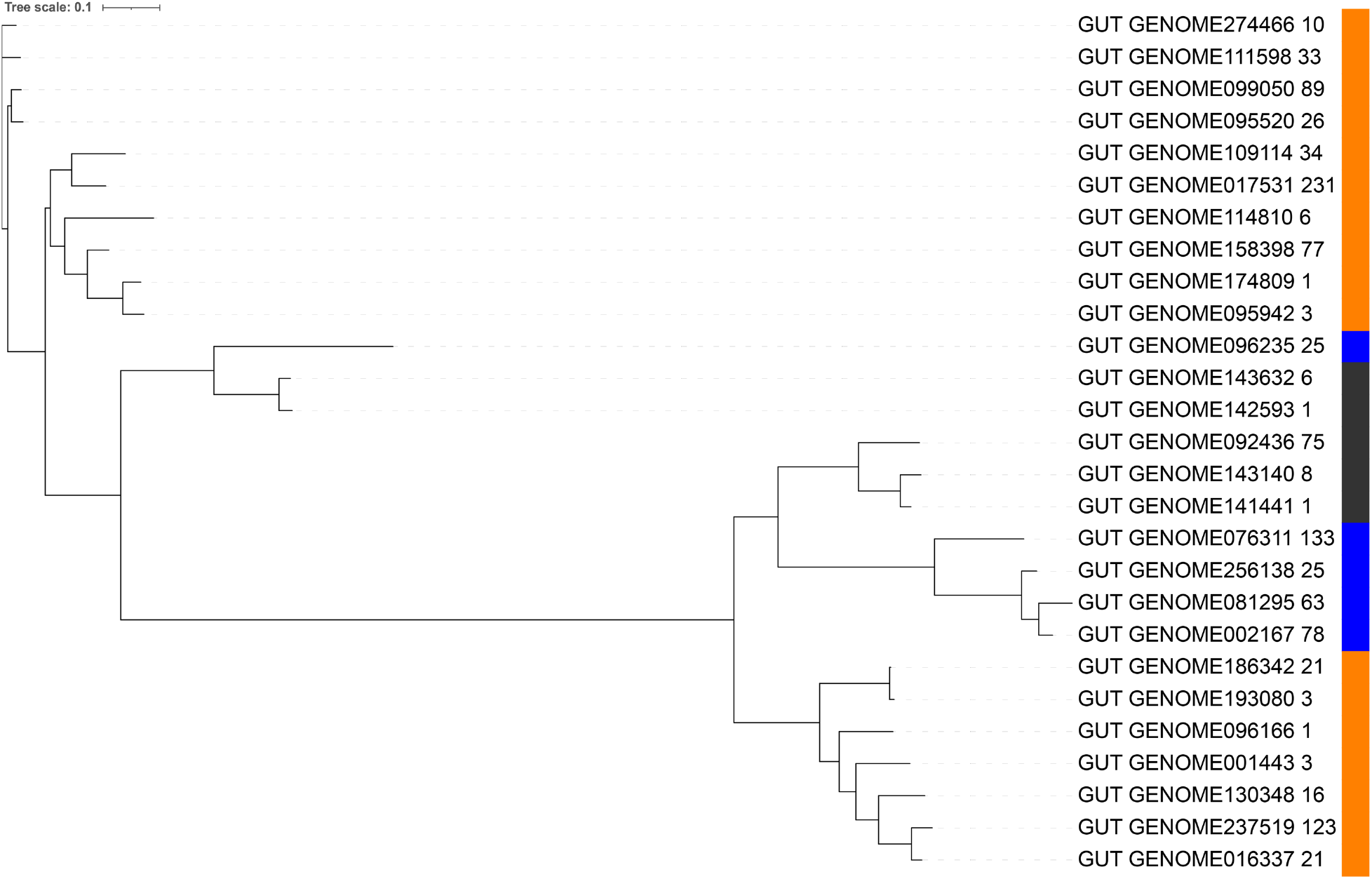
Phylogenetic analysis of dsrA hits from UHGG. Colors on the right signal the phylum that each copy of the gene originated from: orange = *Desulfobacterota_A*, blue = *Firmicutes_B*, black = *Actinobacteriota*. Note: a phylogenetic tree was generated for *dsrB* as well, and it returned an identical tree, therefore, we only included the tree generated from analysis of *dsrA*.

**Supplementary Figure 2.**
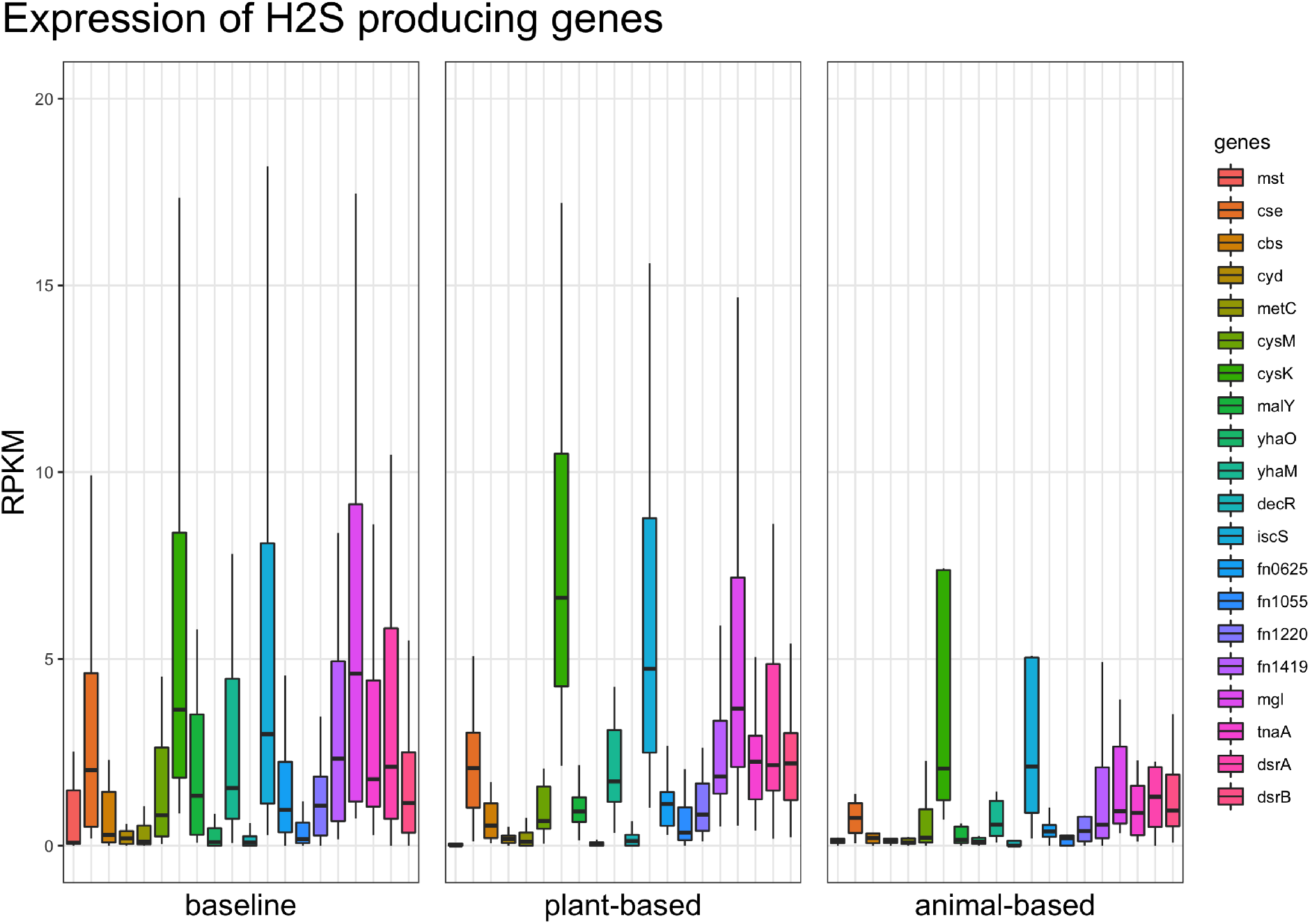
Transcriptomic confirmation for cysteine-degrading genes. This analysis confirms that the H_2_S producing genes considered in this work are actively expressed in healthy humans under a variety of dietary regimens. The y-axis displays log2 transformed RPKM values for each gene (see methods for details on alignment and normalization procedure). The x-axis separates reads counts by both gene and diet intervention. Baseline represents samples taken before either plant- or animal-based diet. Some samples contained zero hits to one or more protein and can be seen along the bottom of the plots. Note that the same individuals were fed the plant and animal based diets with a 6 day waiting period in between the end of one diet and the taking of baseline samples for the next diet.

**Supplementary Figure 3.**
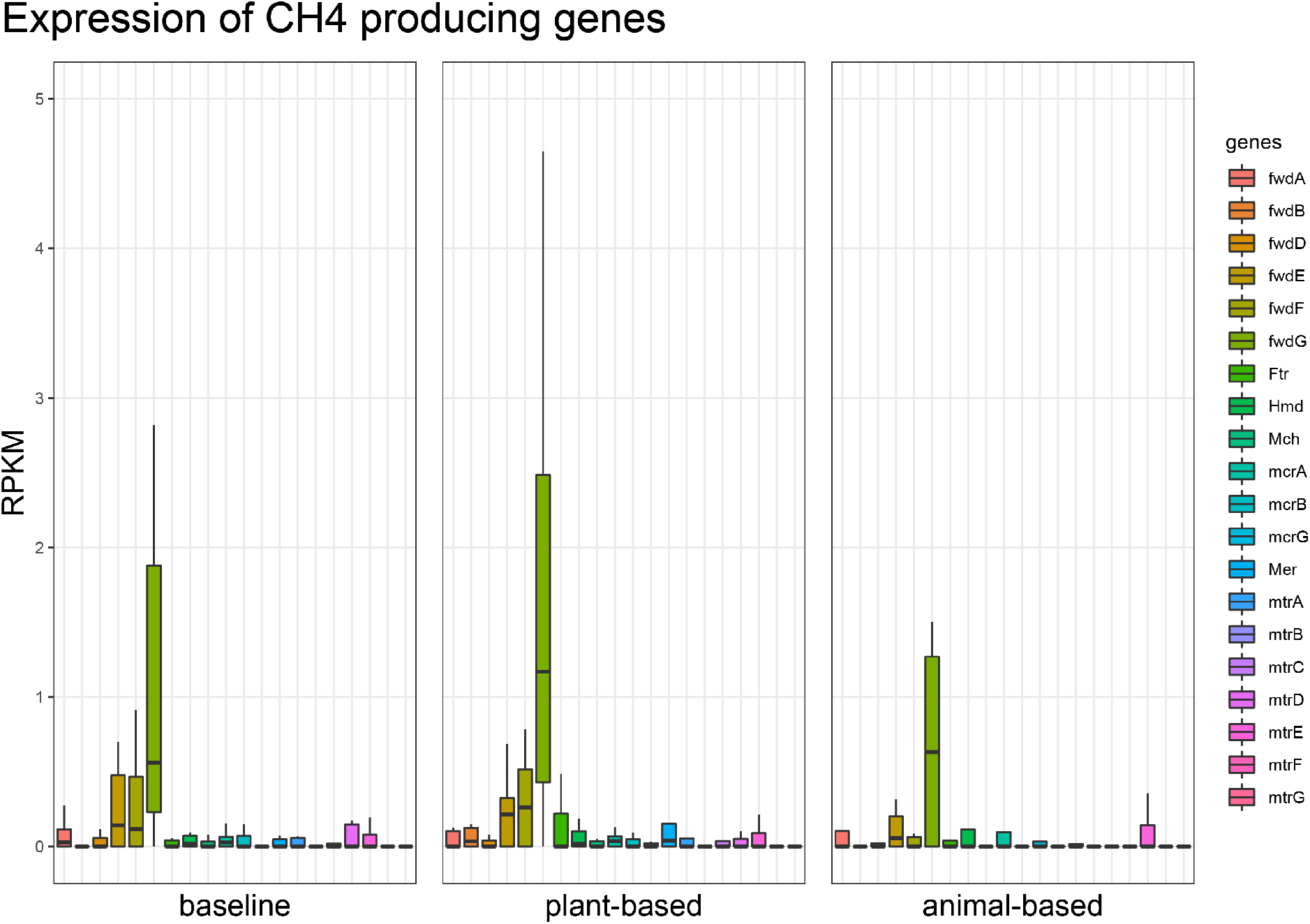
Gene expression of methane producing genes in the human gut. Genes involved in the production of CH_4_ by *Methanobrevibacter smithii* expressed in healthy individuals. The three panels are from 10 participants over three legs of a diet intervention study where metatranscriptomic reads were collected at baseline and diet intervened time points. The y-axis shows RPKM adjusted read counts mapped to each gene involved in the production of CH_4_.

**Supplementary Figure 4.**
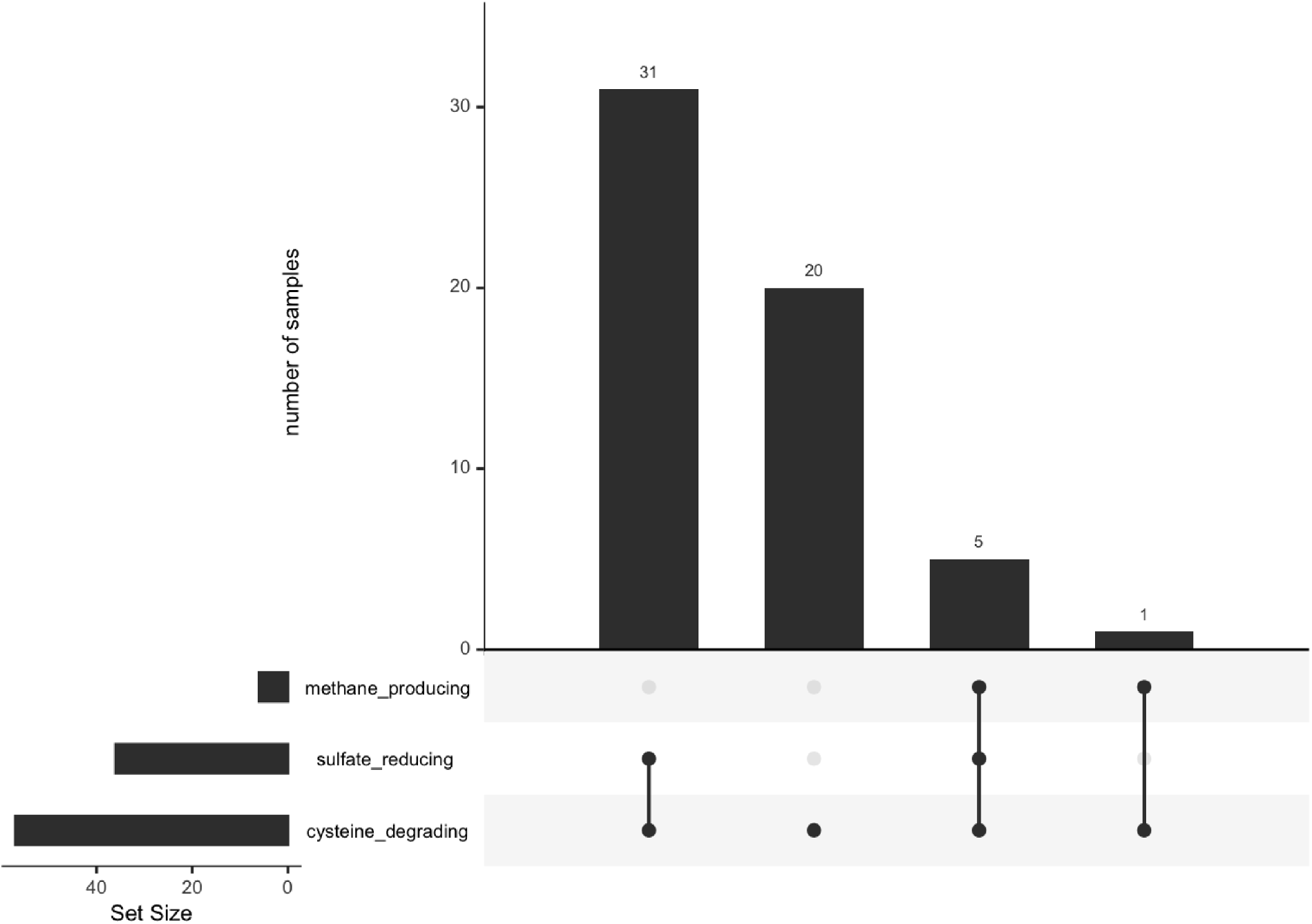
Presence of H_2_S production and CH_4_ production in the healthy human gut microbiome. For some individuals, there is simultaneous production of H_2_S and CH_4_ in the gut microbiome. Out of 59 metagenomic stool samples from David et al. 2014, 57 expressed one or more cysteine-degrading gene, 36 expressed *dsrAB* and 6 expressed at least 90% of the genes required for methane production (see methods). The vertical bars represent the number of samples containing one or more functions described. For example, 31 samples showed expression of sulfate-reducing and cysteine-degrading genes and 5 samples contained the same two functions plus methane production.

