## Supplemental Table 1 caption for "The capacity to produce hydrogen sulfide (H_2_S) via cysteine degradation is ubiquitous in the human gut microbiome"

**Supplementary Table 1. Results of search for putative cysteine-degrading bacteria and sulfate reducing bacteria in UHGG.** A table showing all hits to representative UHGG genome sequences for all $H_2S$ producing genes used in the search space. An aligned gene was considered a hit if the result of the BLASTP search received an E-value < $1x10^{-110}$ and > 50% amino acid identity. The sheet labeled “cysteine_degradation_hits” contains all hits to cysteine-degrading genes as well as curated experimental evidence for $H_2S$ production and details of media used in each reference. The sheet labeled “sulfate_reduction_hits” contains all species receiving hits to dissimilatory sulfite reduction genes *dsrAB*. The sheet labeled “search_space” contains all genes used in the search space to identify putative cysteine-degrading species and sulfate reducing species.
